# Enhanced mRNA delivery using ultrasound-delivered click reactive anchors

**DOI:** 10.1101/2025.01.28.635330

**Authors:** Emilio Di Ianni, Jueun Jeon, Huiyu Hu, Jeremy M. Quintana, Chanseo Lee, Edwina Abou Haidar, Sedra Mohammadi, Ayrton Zargani-Piccardi, Mohammed Mahamdeh, Iván Coto Hernández, Thomas S.C. Ng, Koen Breyne, Hakho Lee, Xandra O. Breakefield, Miles A. Miller

## Abstract

Therapeutic nucleic acid delivery has many potential applications, but it remains challenging to target extrahepatic tissues in a flexible and image-guided manner. To address this issue, we report a bioorthogonal pre-targeting strategy that uses focused ultrasound to promote the delivery of mRNA-loaded lipid nanoparticles (mRNA-LNP). We synthesized amphiphilic click reactive anchors (ACRAs) consisting of a phospholipid PEG-conjugate functionalized with transcyclooctene (TCO) or its companion reactive partner methyltetrazine (mTz), yielding ACRA-TCO and ACRA-mTz. ACRA derivatives were screened for cellular activity, yielding functionalized DOPE-PEG (1,2-dioleoyl-sn-glycero-3-phosphoethanolamine-N- (polyethylene glycol)) derivatives outperforming those containing saturated lipid or branched PEG. Nanobubbles encapsulating ultrasound-responsive gas precursor delivered ACRA-TCO to targeted cells and tissues using focused ultrasound, and this pre-targeting promoted the subsequent delivery of mRNA-LNP functionalized with companion ACRA-mTz. In cell cultures and in mice, ultrasound pre-targeting enhanced the accumulation of mTz-functionalized small molecule and nanoparticle compounds by 75% and 3.6-fold, respectively, and increased gene expression using mRNA-LNP *in vivo*. Taken together, this report presents a modular, ultrasound-enabled strategy for enhancing nucleic acid delivery in targeted tissues.

---

Click-chemistry reactions selectively and efficiently form covalent bonds under physiological conditions, and their ability to perform *in situ* reactions within the body has shown promise for improving localized therapeutic and diagnostic delivery. Various “pre-targeting” strategies have been developed whereby a tissue or cell population is first pre-targeted with an initial agent followed by reaction with a subsequently administered reactive pair^1–3^. Generally, this two-step pre-targeting approach can overcome pharmacokinetic/pharmacodynamic (PK/PD) limitations of traditional one-step strategies. For instance, pairing long-circulating antibodies with short-circulating radionuclides can minimize retention of the latter in blood and off-target tissues^4,5^. Click chemistry reactions also offer potential practical and logistical advantages, since materials like medical radionuclides and ultrasound contrast agents can be short-lived and require special handling^4,6^.

Although click chemistry is not required for pre-targeting, it has proven especially useful as a stable, non-toxic, efficient, and selective means by which to react pairs of reagents in vivo^7^. Click chemistry using trans-cyclooctene (TCO) reacting with methyltetrazine (mTz) through an inverse electron demand Diels-Alder mechanism has an attractive balance of fast reaction kinetics and in vivo stability. Strategies using TCO / tetrazine chemistry have entered clinical testing (NCT04106492).

Various approaches have been developed to pre-target cells and tissues using antibodies^8^, peptides^9^, and small molecules^10^ that engage with molecular targets on the cell surface to present bioorthogonally reactive handles. Yet, known molecular targets are unavailable or undeveloped in some applications due to lack of high expression or expression specificity, ii) desirable internalization/trafficking, iii) safety/tolerability in targeting, and iv) suitable high affinity targeting strategies (ligands, antibodies, and other binders). As an alternative to engaging specific molecular targets on the cell surface, cell surfaces have also been induced to present click-reactive azides by locally delivering azido-sugars to tissue^11^. Despite this progress, a pre-targeting strategy that i) does not depend on specific receptor targeting or cell engineering, ii) can be flexibly targeted for site-specific delivery of cargos (e.g., small molecules, nucleic acids), and iii) promotes cargo internalization and cytoplasmic delivery would be advantageous. Such a strategy would be especially useful for delivering mRNA packaged in lipid nanoparticles (mRNA-LNP), since poor extrahepatic and site-specific delivery remains a barrier to using mRNA-LNP for many biomedical applications^12^.

Here, we report the development of “Amphiphilic Click Reactive Anchors” (ACRAs) to enable site-specific in vivo bioorthogonal reactions with mRNA-LNP. In principle, ACRA-TCO doped into nanobubbles will incorporate into cell membranes upon focused ultrasound treatment, and consequently promote the cellular binding and delivery of a subsequently administered mTz-functionalied mRNA-LNP.

To begin developing this approach, we first synthesized a targeted panel of TCO-functionalized lipid-PEG conjugates (ACRA-TCO) (Figure 1A, S1, S2) and measured their ability to present reactive TCO moieties on the surface of two different live cell cultures (Figure 1B). Using the murine YUMM1.7 melanoma cell line or the MOC2 (Mouse Oral Squamous Carcinoma) cell line, we treated cells for 30 min. with the panel of ACRA-TCO conjugates at 15 μM in complete cell culture medium, rinsed twice with phospho-buffered saline (PBS), treated cells with 15 μM mTz-AlexaFluor488 (mTz-AF488) and incubated for 10 min. We then rinsed cultures twice and immediately imaged cells for fluorescence accumulation. Negligible fluorophore accumulation occurred in the absence of ACRA-TCO treatment. All ACRA-TCO derivates yielded detectable mTz-AF488 accumulation, with DOPE-PEG_2k_-TCO (**3;** 1,2-dioleoyl-sn-glycero-3- phosphoethanolamine-N-(polyethylene glycol)_2k_-TCO) showing the highest fluorescence in both cell lines, followed by DOPE-PEG_1k_-TCO (**2**; Figure 1B). The unsaturated lipid-conjugate DOPE-PEG_1k_-TCO (**2**) showed a 2.6±0.2-fold higher signal than the saturated lipid equivalent, DSPE-PEG_1k_-TCO (**1;** 1,2- Distearoyl-sn-glycero-3-phosphoethanolamine-N-(polyethylene glycol)_1k_-TCO), suggesting the former more efficiently inserts into the plasma membrane’s phospholipid bilayer. ACRA with branched PEG structures and multivalent TCO failed to outperform the monofunctional single PEG chain structures despite the 3-fold molar excess of reactive TCO. Based on a calibration curve with mTz-AF488, and averaged dye captured on single cell, there was an equivalent of 0.4 µM / cell from applied 2µM / cell, resulting in a roughly 20% click reaction yield and 240,000 molecules/cell reacting on the cell surface with mTz-AF488. Taken together, ACRA-TCO with monovalent PEG-TCO and an unsaturated DOPE (1,2- dioleoyl-sn-glycero-3-phosphoethanolamine) dilipid anchor showed ideal membrane incorporation and TCO reactivity, allowing efficient bioorthogonal capture of mTz-Dye.

**Figure 1.**
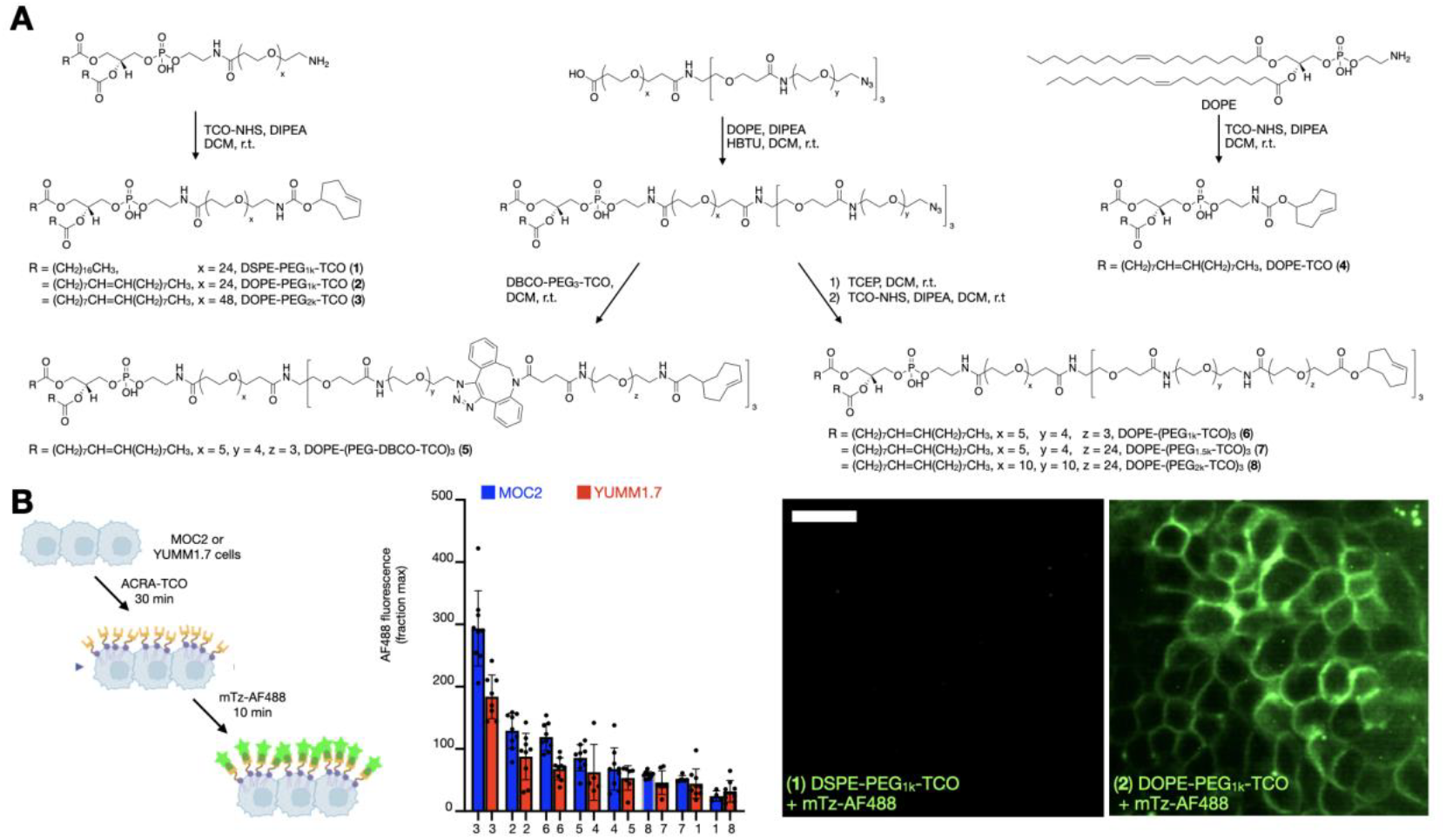
“Amphiphilic click reactive anchors” (ACRAs) label cell membranes with a click reactive moiety. **A**) Synthetic scheme of a panel of ACRA-TCO conjugates **1**-**8. B)** Experimental schematic (left), quantification (middle), and representative fluorescence microscopy images (right, MOC2 cells) evaluating the ability of various ACRA-TCO to incorporate into MOC2 or YUMM1.7 cultures. Scale bar, 50 µm. Data represents mean ± SD, n=3.

We next tested the kinetics of ACRA incorporation into live cell membranes. We used the top-performing ACRA-TCO compound **3** (DOPE-PEG_2k_-TCO) on MOC2 cells. Cells were treated with ACRA-TCO for between 5 min and 1 hr, washed to remove unbound material, treated with the fluorophore mTz-AF488 at equimolar 20 μM, washed, and immediately imaged (Fig. 2A). Labeling was detectable by 5 min and nearly saturated by 30 min. Fitting to a saturable binding model indicated half-maximum incorporation approximately 11 minutes; 25% of cells showed detectable ACRA incorporation by 5 minutes, which increased to >95% after 1 hr of labeling. Since mTz-AF488 is negatively charged and impermeable to cell membranes, fluorescence predominately arises from extracellular reactions on the cell surface, and this is reflected by the membrane-staining pattern in the images. Time-lapse microscopy in a controlled 37^°^C and 5% CO_2_ environment showed that cells were viable and retained lipid-PEG conjugate for at least 16 hrs following DOPE-PEG-AF488 treatment and washing (**Supplement Video 1**). Given the bimolecular kinetic constant reported for TCO/mTz coupling with bioconjugates (k_act_ ∼ 500 M^-1^ s^-1^)^7,13^, the estimated concentration of ACRA on the cell surface (∼0.5 μM), and mTz-AF488 in solution (20 μM), the theoretical TCO/mTz coupling would reach a half-maximum in ∼1 minute and reach >95% completion by 10 min. Therefore, the availability of functional TCO on the cell surface is the limiting factor in the *in vitro* experiments, rather than TCO/mTz conjugation kinetics. Overall, these data show ACRA incorporates into cell membranes within minutes and presents reactive TCO on the cell surface for at least 60 min, a time window amenable for pre-targeting in vivo.

**Figure 2.**
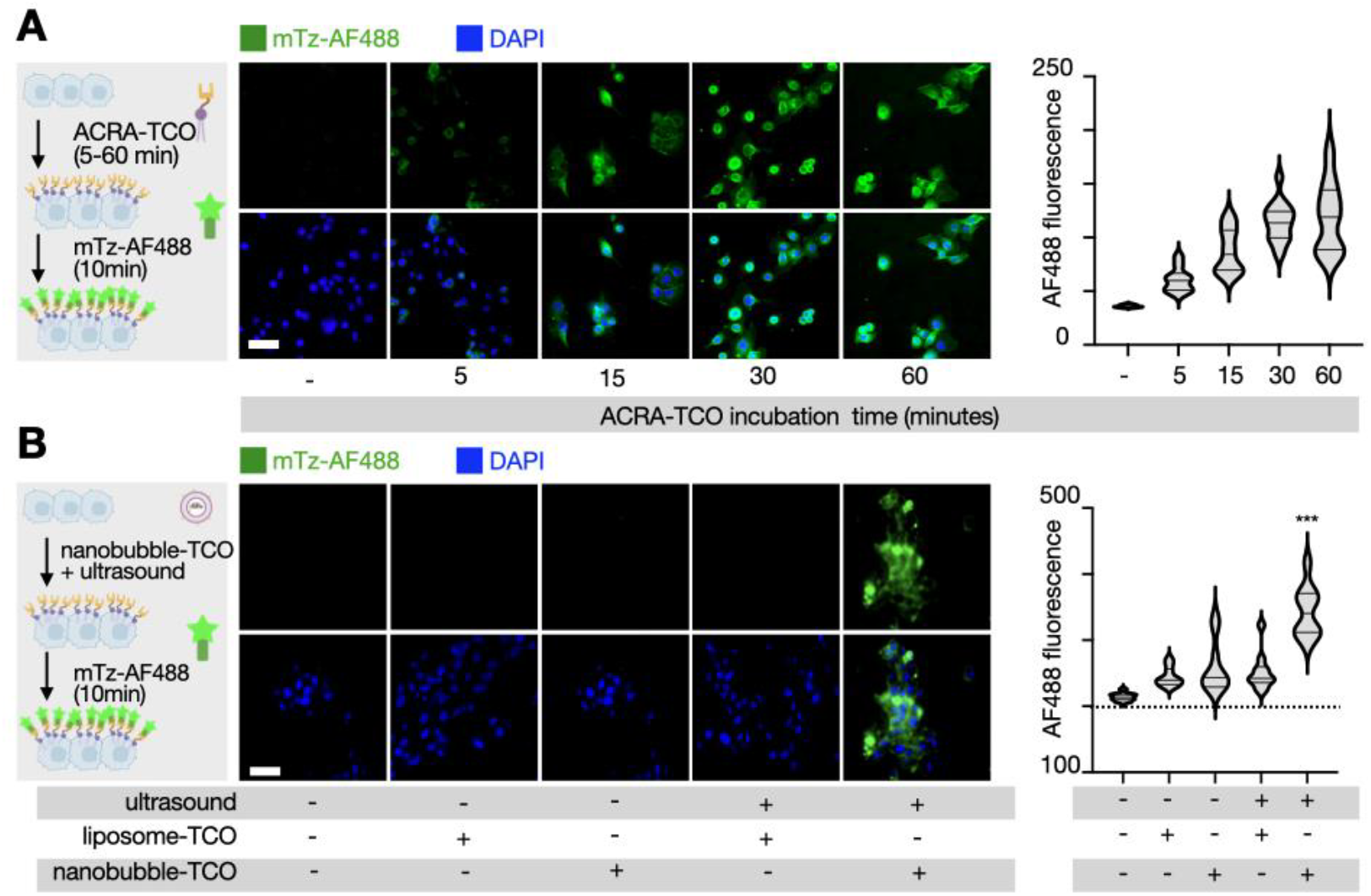
Ultrasound-responsive nanobubbles control the rapid incorporation of ACRA into cell membranes. **A)** Experimental schematic (left), representative images (middle), and quantification (right) of ACRA-TCO membrane labeling kinetics in MOC2 cells. **B)** Experimental schematic (left), representative images (middle), and quantification (right) of YUMM1.7 cells treated with ultrasound-treated ACRA-doped nanobubbles. Scale bar, 100 µm. Data are median and quartiles. \*\*\**P* < 0.001, n=3.

With evidence that free ACRA efficiently incorporates into cell membranes, we investigated whether this behavior could be controlled by ultrasound. We first produced nanobubbles (ACRA-free) and characterized their properties including size and bursting efficiency, when sonicated. For this purpose, ACRA-free nanobubbles were prepared by combining DSPC (1,2-distearoyl-sn-glycero-3-phosphocholine), cholesterol, and DSPE-PEG_2k_ dissolved in chloroform at a molar ratio of 67.4:27.6:3 mol% to obtain a lipid film, which was then hydrated to obtain a suspension of liposomes. These were processed by serial lipid extrusion through membrane filters with pore sizes of 400 nm and 50 nm. A probe-type sonicator then encapsulated gas precursor (perfluorohexane, PFH) into the liposomes. After PFH encapsulation, a slight increase in overall size was observed, from 84 nm to 164 nm in hydrodynamic diameter (D_h_) as measured by dynamic light scattering (DLS), with a final polydispersity index (PDI) of 0.23 (Figure S4, center). The stability of the nanobubbles was maintained for 36 hrs at 37 °C PBS (Figure S4, right). We applied ultrasound to these nanobubbles and observed strong signal contrast under an ultrasound-imaging mode (Figure S5). Ultrasound imaging revealed cavitation-induced nanobubble disruption (“popping”) under acoustic powers from 5W to 20W and across duty cycles of 20% and 50%. To balance nanobubble popping efficiency with potential cell toxicity, ideal ultrasound popping parameters were chosen at 5W with a 50% duty cycle. Negligible impacts on cell viability were observed under this condition (Figure S5).

We next functionalized nanobubbles with ACRA-TCO, and evaluated two candidates: the single-chained DOPE-PEG_1k_-TCO (**2**) and the branched PEG construct DOPE-(PEG_1k_-TCO)_3_ (**6**). Nanobubbles incorporated either 1% or 2% molar ratio of each ACRA, and DLS indicated that ACRA-doped nanobubbles maintained similar size ranges (D_h_ between 140 and 160 nm) regardless of the formulation (Figure S6). When reacted with the fluorophore mTz-Cy5, ACRA-doped nanobubbles (2) yielded fluorescent signals, validating that the TCO moiety in the ACRA nanobubbles remained reactive (Figure S6). However, we observed variations in the fraction (*r*) of clicked TCO over TCO loading. The value was the maximal (*r* = 90%) with nanobubbles containing 2% single-armed ACRA, followed by nanobubbles containing 1% single-armed ACRA (**2**) (*r* = 86%). Nanobubbles with three-armed branched ACRA **(6**) were less effective in reacting with mTz-Cy5 (*r* = 63% for 1% three-armed ACRA; *r* = 61% for 2% three-armed ACRA), presumably due to steric hindrance between closely spaced TCO molecules. Based on these results, we opted to use a formulation of a 2% molar ratio of single-armed ACRA **(3)**. Fluorescence quantification of the Cy5-labeled nanobubbles confirmed that adding ACRA **(3)** to the nanobubble formulation did not substantially affect ultrasound bursting responsiveness (Figure S7).

Given these results, we examined whether bursting ACRA-doped nanobubbles with ultrasound could selectively allow ACRA to incorporate into cell membranes (Figure 2B). Live YUMM1.7 cells were treated with either native nanobubbles lacking ACRA, ACRA-doped nanobubbles, or ACRA-doped liposomes lacking gas precursor, which were exposed to ultrasound or not (Figure 2B). Following 30 min treatment with these materials, cells were rinsed and treated with mTz-AF488, washed, and imaged by microscopy to quantify the amount of fluorescent mTz-AF488 that had been able to react with TCO presented on the cells. Results showed that only ultrasound-treated ACRA-doped nanobubbles enhanced mTz-AF488 staining, and fluorescence subcellular localization was consistent with cell surface localization (Figure 2B). Taken together, these results indicate ultrasound bursting of nanobubbles liberates ACRA dopants and promotes their incorporation into live cell membranes within minutes.

Next, we assessed whether the pre-targeting with ACRA-doped nanobubbles could improve the ability of cells to accumulate LNPs. We hypothesized that ACRA-TCO on cell membranes could covalently bind mTz-doped LNPs, thereby holding them to the cell surface and promoting their cellular uptake. To begin testing this hypothesis, we first quantified cellular accumulation of fluorophore-doped LNPs by fluorescence microscopy. Guided by prior development^14^, LNPs were formulated with the ionizable lipid SM-102, β-sitosterol, helper lipid DOPC (1,2-dioleoyl-sn-glycero-3-phosphocholine), and the PEG2000 DMG (1,2-dimyristoyl-rac-glycero-3-methoxypolyethylene glycol-2000) (Figure S8). In addition, a lipid conjugate of the fluorophore Cy5 (N-palmitoyl sphingomyelin-N-(Cyanine 5), “C_16_-Cy5”) was doped into the lipid mixture during the initial formulation. Dynamic light scattering showed hydrodynamic diameter (D_H_) and PDI of 102 nm and 0.22, respectively. We then functionalized LNPs by incubating them with ACRA-mTz for 2 hrs and purifying them by a 30 kDa MWCO (molecular weight cut-off) ultracentrifugation filter to remove unincorporated free ACRA. DLS showed only slight increase in size with ACRA doping (Figure S9; D_H_ 110 nm, PDI 0.25). We verified mTz functionalization of LNP by incubating mTz-LNPs with TCO-AF488 and measuring reaction products by fluorescence microscopy. Results showed particulate colocalization between C_16_-Cy5, which was doped into LNPs during their formulation as above, and AF488, therefore indicating successful conjugation between mTz-LNP and TCO- AF488. A degree-of-labeling analysis indicated roughly 6 functional mTz molecules per individual LNP, which was calculated by AF488 fluorimetry combined with nanoparticle tracking analysis to measure LNP concentration (Figure S9).

Having demonstrated mTz LNP functionalization, we next tested whether ultrasound-mediated ACRA pre-targeting could promote LNP accumulation in live cells (Figure 3). Nanobubbles were incubated with YUMM1.7 cells for 1 hr with or without ultrasound treatment, cells were washed, and then treated with mTz-LNP. Under these conditions, fluorescence microscopy showed LNP uptake only in cells exposed to ultrasound-treated nanobubbles with 8 to 30-fold higher uptake in Cy5 fluorescence compared to all other treatments (Figure 3). mTz-LNP showed distinct subcellular distribution compared to mTz-AF488 when added to cells pre-targeted with ACRA-TCO (Figure 3). Specifically, after reacting with ACRA-TCO on the membrane, mTz-AF488 appeared localized on the cell membrane after 1hr, while mTz-LNPs were already internalized into the cells. These data suggest that ACRA-TCO facilitates cellular capture of LNP while permitting their internalization, which is a necessary step of delivering mRNA to the cytoplasm.

**Figure 3.**
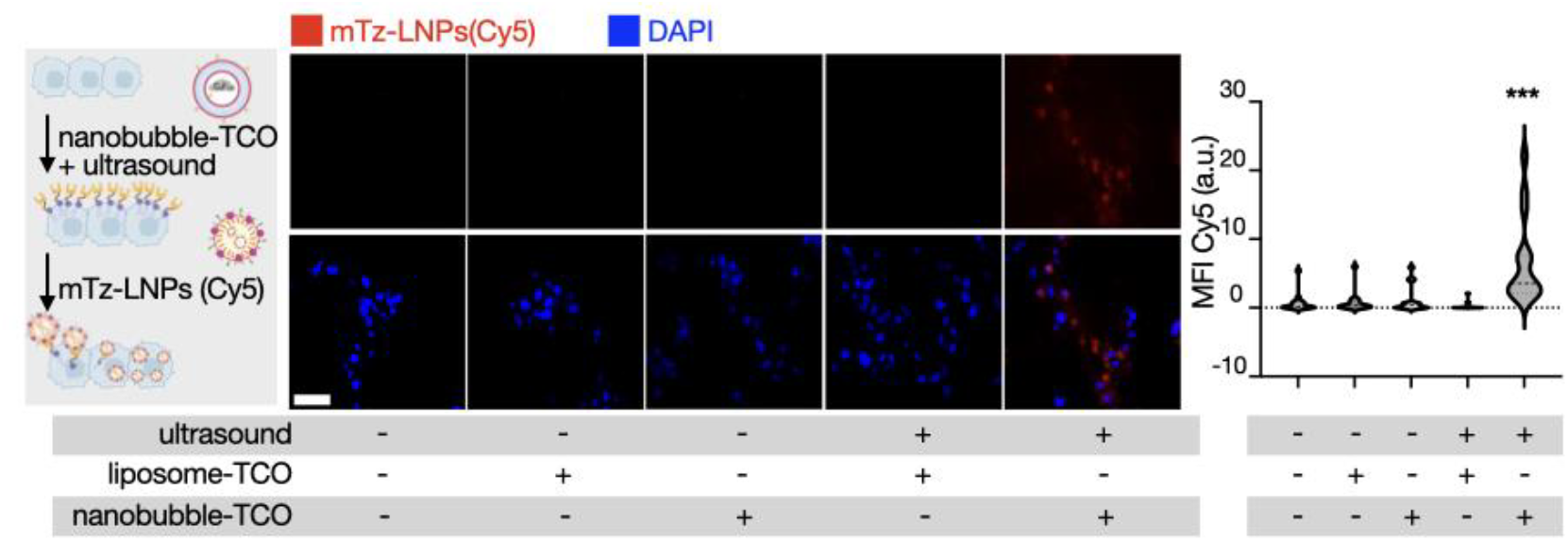
ACRA-doped nanobubbles enhance cellular LNP delivery. Experimental schematic (left), representative fluorescence microscopy (center), and quantification of Cy5 MFI (mean fluorescence intensity, right). Scale bar, 100 µm. Data are median and quartiles. \*\*\**P* < 0.001 and n=3.

We next tested the degree to which ultrasound could combine with ACRA-doped nanobubbles to enable site-specific pre-targeting of LNP in mouse tissue (Figure 4A). We used Cy5-labeled mTz-LNP or free mTz-Cy5 fluorophore, with the latter used as a model small molecule compound (Figure 4B). An initial nanobubble circulating half-life (*t*_*1/2*, *initial*_) of 14 min was determined from fluorimetry of blood samples collected following intravenous injection of Cy5-labeled nanobubbles into healthy C57Bl/6J mice (Figure S11), therefore confirming that nanobubbles circulate long enough to respond to ultrasound treatment within minutes following their administration. To assess pre-targeting abilities, mice were first intravenously injected with ACRA-doped nanobubbles followed by ultrasound (5 W, 50% duty cycle, and 1 min) to the hindleg skin immediately after. Cy5-labeled mTz-LNP or free mTz-Cy5 dye was injected intravenously roughly 3 min later. Tissues were harvested 3 hrs later and imaged for fluorescence (Azure Sapphire FL). Results with mTz-LNP showed 3.6 ± 0.9-fold enhanced fluorescence in ultrasound-treated skin compared to untreated skin on the contralateral hindleg (Figure 4B and S12). As expected, the free mTz-Cy5 dye accumulated in skin even without ultrasound, but signals were still 75% ± 28 higher in treated compared to untreated tissue. Thus, ultrasound promoted the delivery of both LNP and a small molecule fluorophore as two model agents with highly distinct physicochemical and pharmacokinetic properties.

**Figure 4.**
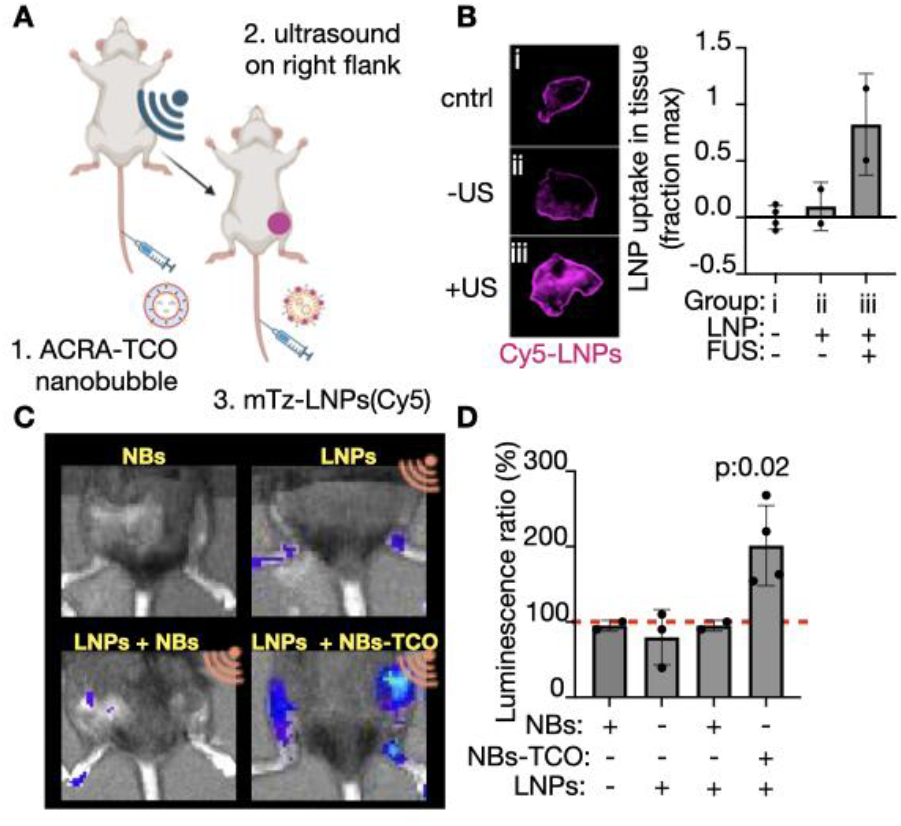
Ultrasound-activation enhances LNP delivery and mRNA expression in mouse tissue. A) Experimental schematic. B) Representative Cy5 fluorescence imaging and corresponding quantification indicating local increase of dye delivered by LNPs in the target pretreated with CRA TCO nanobubbles and ultrasound (US). C) Representative bioluminescence imaging and D) corresponding quantification indicating local expression of luciferase mRNA delivered by LNPs coupled with ACRA-TCO nanobubbles and US and controls. Data are means +/− std. dev.

Since we found that ACRA-doped nanobubbles could increase the accumulation of LNP in ultrasound-treated tissue, we next investigated whether this could translate into increased tissue expression of mRNA delivered by such LNP. As a model reporter system, we used luciferase mRNA (N1-methylpseudouridine base-modified FLuc), since its expression is quantifiable by bioluminescence imaging following administration of a D-luciferin substrate. To specifically evaluate the functional impact of *in vivo* TCO/mTz conjugation, control groups in this experiment also included native nanobubbles lacking ACRA-TCO. Similar to the experiment above, mice were first treated with nanobubbles, followed by ultrasound to the right hindleg immediately after (5W, 50% duty cycle, 1.2W/cm2, 1MHz, with 30s on and 10s off, 3 min. exposure), immediately followed by intravenous mRNA-LNP right after. Live bioluminescence imaging was performed 4 hrs later to quantify luciferase expression (Figure 4C). Results showed that local ultrasound treatment improved luciferase levels only when combined with mTz-LNP and ACRA-doped nanobubbles, as evidenced by a 2-fold enhanced luciferase expression in the right hindleg of mice that were treated with ACRA-doped nanobubbles, ultrasound and LNP (Figure 4D). Nanobubbles lacking ACRA were ineffective at increasing luciferase expression, therefore indicating that *in vivo* TCO/mTz conjugation contributed to enhanced mRNA delivery. The experiments with mRNA delivery showed hepatic delivery by bioluminescence imaging (Figure S13). This result is not unexpected, since LNP similar to the formulation used herein avidly accumulate in the liver, and the stoichiometry is such that ultrasound-targeted delivery is not acting as a systemic sink of LNP that prevents them from acting elsewhere in the body. Nonetheless, these results establish a strategy for using focused ultrasound to improve the local delivery and expression of mRNA in a targeted and flexible manner.

This report presents a strategy for using ultrasound and *in vivo* bioorthogonal chemistry to pre-target cells and improve mRNA-LNP delivery. Prior work has described *in vivo* bioorthogonal chemistry for enhanced molecular imaging, radiopharmaceutical therapy, chemotherapy, antimicrobials or prodrug delivery, yet a need still exists for strategies that allow site-specific delivery of mRNA^12^. The strategy presented herein builds on prior work and offers several key advantages. We identify DOPE-PEG-TCO (**2**-**3**) and DOPE-PEG-mTz as amphiphilic lipid-PEG conjugates that can stably insert into biological membranes and LNPs, respectively. DOPE-PEG-TCO exhibits superior membrane labeling kinetics compared to related lipid- PEG conjugates including DSPE-PEG-TCO and those containing branched PEG, with the latter being likely hindered by sterics. DSPE-PEG has saturated fatty acid chains, leading to a high phase transition temperature, tight and rigid packing, and therefore slower membrane insertion kinetics compared to DOPE-PEG. DSPE-PEG and other unsaturated lipids have been extensively used in past work to anchor materials to the cell surface^15^, and DOPE-PEG-TCO may offer advantages in other such contexts such as delivery of a click reactive compounds by ultrasound and click-doped nanobubbles. We show that DOPE-PEG-TCO incorporates into nanobubbles, which controls site specificity of bio-orthogonal reactions.

Limitations of the study are noted. We focused on enhancing delivery to ultrasound-treated tissue rather than minimizing off-target activity. Future work may refine the strategy to minimize off-target accumulation, while increasing the likelihood to react with ACRA on the target site. In this line, potential modifications include altering the LNP formulation to promote charge-dependent tropism^16^ or increasing the circulation time of LNPs^17^, so to increase chances of interactions between circulating LNPs and click handles on the target site. Ultrasound itself may affect drug delivery by permeabilizing vasculature and allowing macromolecular vehicles such as mRNA-LNP to extravasate from the blood into tissue interstitium^18^. Nonetheless, in this work we found that such non-specific effects did not impact mRNA-LNP delivery in the absence of TCO/mTz conjugation. Thus, ACRA-doped nanobubbles present a flexible, image-guided strategy for targeting tissue for delivery of mRNA-LNP and other relevant agents. This method holds promise for mRNA therapies directed at difficult-to-reach tissues lacking ideal receptors for traditional molecular targeting.

## Acknowledgments

This work was supported by Ionis Pharmaceuticals. XOB is funded by NCI CA232103. KB is funded by NIH K22 CA2802019-01 and DoD HT9425-24-1-0119. J.Q. was supported in part by US National Institute of Health grants DP2CA259675 and DP2CA259675-01S1 (M.A.M). Non-financial support from Genentech/Roche, Pfizer, and Archon Biosciences (M.A.M) and Bayer (T.N.) is acknowledged, all outside the submitted work. TSCN is supported by R21EB036323, R01DK097112, R01EB034692, the U.S. Department of Defense(W81XWH-22-1-0061), and the Lee Family Foundation, and receives unrelated research support from Bayer.

